# Using light for energy: examining the evolution of phototrophic metabolism through synthetic construction

**DOI:** 10.1101/2022.12.06.519405

**Authors:** Autumn Peterson, Carina Baskett, William C. Ratcliff, Anthony Burnetti

## Abstract

The origin of phototrophy was pivotal in increasing the size and scale of the biosphere, as it allowed organisms to utilize light-driven energy transport to drive biological processes. Retinalophototrophy, one of two independently evolved phototrophic pathways, consists of a simple system of microbial rhodopsins which have spread broadly through the tree of life via horizontal gene transfer. Here, we sought to determine whether *Saccharomyces cerevisiae*, a heterotrophic fungus with no known evolutionary history of phototrophy, can function as a facultative artificial phototroph after acquiring a single rhodopsin gene. We transformed *S. cerevisiae* into a facultative phototroph by inserting a rhodopsin protein from *Ustilago maydis* into the yeast vacuole, allowing light to pump protons into the vacuolar compartment, a function typically driven by consuming ATP. We show that yeast with rhodopsins gain a selective advantage when grown under green light, growing more rapidly than their non-phototrophic ancestor or rhodopsin-bearing yeast cultured in the dark. These results underscore the remarkable ease with which rhodopsins may be horizontally transferred even in eukaryotes, providing novel biological function without first requiring evolutionary optimization.

## Introduction

Major evolutionary innovations have shaped the history of life on Earth. Some major innovations occurred multiple times: for example, at least 50 distinct multicellular lineages have arisen from a diverse range of unicellular ancestors[1, 2]. Other transitions, such as eukaryogenesis, have evolved only once [3]. One of the most impactful innovations in evolutionary history is phototrophy – the ability to use light for biological energy. Originating at least 3.5 billion years ago[4], this process drives the vast majority of metabolism on Earth, directly or indirectly[5]. The ability to utilize light for biological energy has evolved twice independently[6], with a complex history involving both vertical and horizontal transfer.

There are two types of phototrophic metabolism: chlorophototrophy and retinalophototrophy. Chlorophototrophy, observed only in a subset of bacteria including the oxygenic cyanobacteria and numerous anoxygenic phototrophs[7], uses photochemical reactions in chlorophyll and bacteriochlorophyll molecules to excite electrons to reducing potentials and either fix carbon or drive an electron transport chain to pump protons. This system is complex, with long metabolic pathways required to construct chlorophyll pigments, large membrane complexes with multiple components transducing energy, and multiple redox cofactors[5]. In contrast, retinalophototrophy utilizes a much simpler system consisting of a single microbial rhodopsin protein[8]. This small membrane protein is covalently bound to all-trans-retinal pigment and pumps a single proton per photon absorbed rather than engaging in light-driven redox reactions[9].

Microbial rhodopsins can be found in all three domains of life[10, 11], unlike chlorophototrophic reaction centers, which are only found in eubacteria or eukaryotes that have taken up endosymbionts[7]. This difference is thought to be due to the ease with which rhodopsins can be transferred horizontally, as they are encoded by a single gene[8] with only four additional genes required to synthesize the necessary retinal pigment cofactor[9, 12, 13]. In contrast, at least thirty genes are required for horizontal transfer of a chlorophototrophic pathway[14]. Horizontal transfer of chlorophototrophy in the deep past has been inferred[14, 15], but is rare compared to the phylogenetic ubiquity of rhodopsins. The simplicity and ease of transfer of microbial rhodopsins, along with experiments regarding the physiological effects of this protein in encouraging recovery from starvation, suggests that they provide an evolutionarily accessible alternative phototrophic metabolism that is beneficial in resource-limited environments[16, 17]. As an alternate source of energy that does not depend on consuming biomass, they represent a way to provide a basic energy metabolism without consuming scarce resources or to extend quiescence beyond what would ordinarily be possible.

The comparative record leaves little doubt that rhodopsins are easily transferred horizontally[11, 16, 18–22]. However, little is known about the dynamics through which a heterotroph can be transformed into a nascent phototroph. While there are multiple reported experiments demonstrating heterologous expression of rhodopsins in *E. coli* that provide a selective advantage[23–26], it is sometimes unclear whether the selective advantage is due to phototrophy or changes in membrane structure[23]. No similar experiments have been performed in a eukaryotic host cell, with rhodopsins from another species that have not been modified. Bacteria and archaea may be pre-adapted for using rhodopsins for biological energy, as they use a single cellular membrane for both structural and bioenergetic purposes. Protons pumped across this membrane via rhodopsins readily translate into ATP, and newly introduced membrane proteins readily localize to this same membrane. Eukaryotes typically use oxidative phosphorylation on the inner mitochondrial membrane for their energy metabolism, which is significantly more complex in terms of protein trafficking mechanisms and protein specialization compared to doing so on bacterial membranes[27]. Can eukaryotes with no evolutionary history of phototrophy use newly acquired rhodopsins to generate biological energy with a single horizontal gene transfer event, or does this process require more extensive evolutionary tuning than in prokaryotes?

Here, we attempt to transform *Saccharomyces cerevisiae*, a heterotrophic fungus with no known evolutionary history of phototrophy, into a facultative artificial phototroph by introducing a fungal rhodopsin. We transferred a rhodopsin that localizes to the vacuole, an organelle which is normally acidified via a rotary proton-pumping V-type ATPase, thus allowing the rhodopsin’s light-driven proton pump to supplement heterotrophic metabolism. Probes of the physiology of modified cells show that they are able to deacidify the cytoplasm using light energy, demonstrating the ability of rhodopsins to ameliorate the effects of starvation and quiescence. Rhodopsin-bearing yeast show a selective advantage in the presence of green light, which is due to increased growth, not reduced death. Together, our results highlight the remarkable ease with which an obligate heterotroph, *S. cerevisiae*, can be transformed into a facultative photoheterotroph.

## Results

### Strain construction/validation

We chose the vacuolar rhodopsin gene *UmOps2* from the fungus *Ustilago maydis*[28] to introduce into unicellular *Saccharomyces cerevisiae* (strain Y55). This rhodopsin localizes to the vacuole membrane, one of two membranes in a eukaryotic cell containing a rotary ATPase theoretically capable of transducing a proton gradient into ATP (the other being the inner mitochondrial membrane)[29] Attempts to introduce a proteorhodopsin directly from bacteria[30] resulted in localization to the endoplasmic reticulum rather than either of these membranes. While rhodopsins have been synthetically engineered to localize to the inner mitochondrial membrane of eukaryotes via fusions to mitochondrial inner membrane proteins[31–34], this has never been observed in nature and thus would be a less suitable model for a natural horizontal gene transfer event.

*UmOps2* and *UmOps2-GFP* fusion genes were synthesized and codon optimized for yeast, and introduced to the yeast chromosome via a plasmid bearing homology to the HO gene locus (Figure 1A). Microscopy indicated that localization to the vacuolar compartment, previously observed in *U. maydis*, was maintained (Figure 1B). This rhodopsin is oriented to pump protons into the vacuolar compartment when energized with light, a process that may either build up the vacuolar membrane potential sufficiently to run the vacuolar ATPase in reverse and generate ATP, or simply reduce ATP consumption by the vacuolar ATPase, an enzyme that incurs a significant fraction of the cellular ATP budget[35] (Figure 1C). To absorb light, rhodopsin must be covalently bound to retinal, a chromophore that converts light into metabolic energy by converting one bond from trans to cis, with this shape change forcing a proton across the membrane[8, 36]. We observed that yeast bore a purple color when all-trans-retinal is added to cells containing rhodopsin, indicating that the apoprotein is able to successfully integrate this chromophore into the complex (Figure 1D).

**Figure 1.**
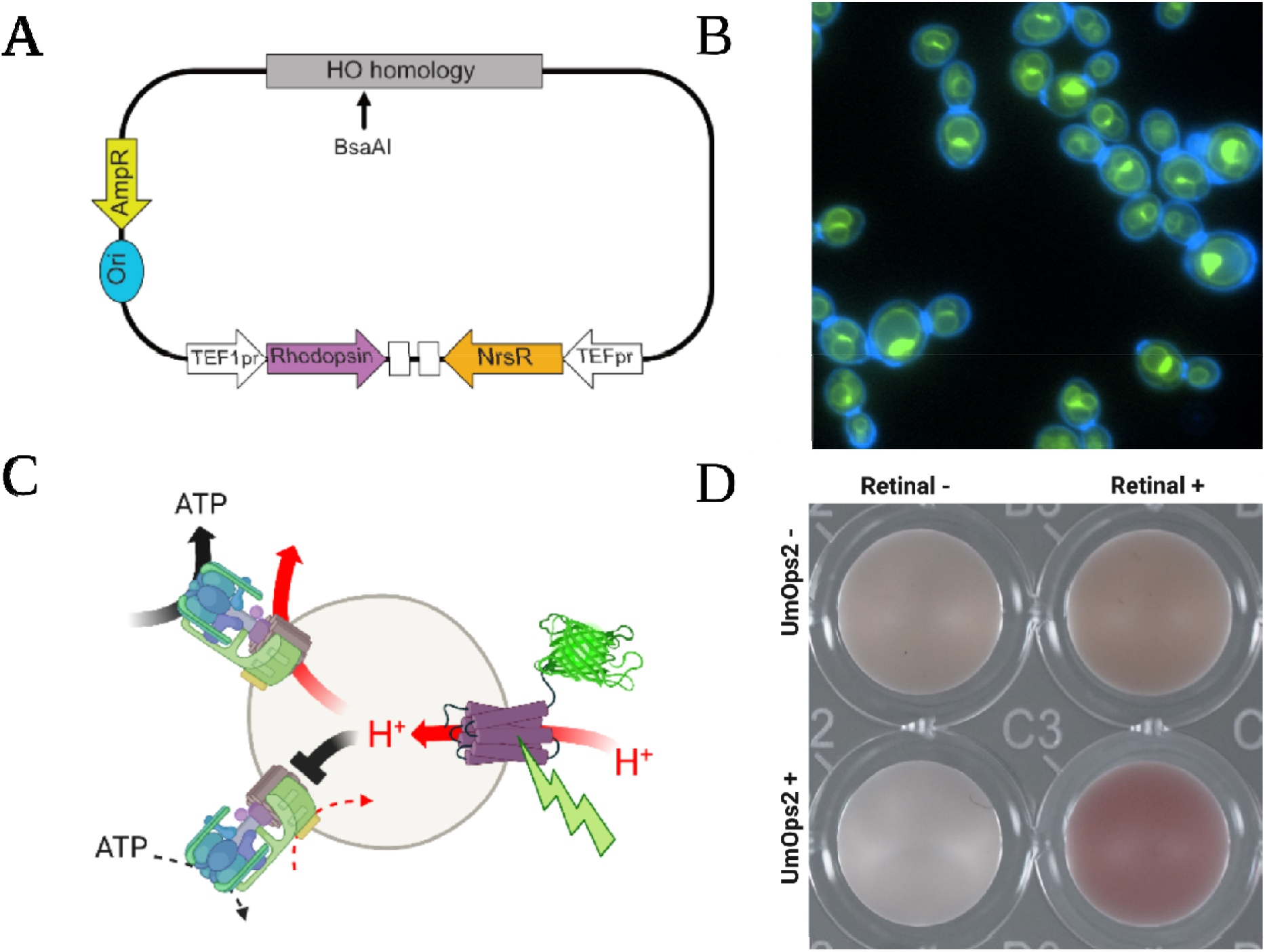
Constructing rhodopsin-bearing yeast. (A) *UmOps2* rhodopsin was put under the control of the *TEF1* promoter, on a plasmid with the *NatMX6* selectable marker and a segment of homology to the *HO* gene of yeast. This homology region was cut with restriction enzyme BsaAI for ends-in cloning to produce a multicopy repeat array at the HO locus. (B) The UmOps2-GFP fusion protein localizes to the vacuolar membrane (cell walls stained with calcofluor in blue). (C) In the presence of green light, rhodopsin pumps protons from the cytoplasm into the vacuolar compartment. This either reduces ATP consumption by the rotary ATPase, or pushes the vacuolar membrane to a high enough voltage to allow ATP production rather than consumption. (D) Confirmation of UmOps2 protein expression and function. Yeast bearing the transgenic rhodopsin gene exhibit a purple color, but only when both the transgene is present and the yeast are supplemented with all-trans retinal.

### Physiological analysis

To determine if rhodopsins affect the physiology of *S. cerevisiae*, we measured the effects of green light on yeast with and without UmOps2 on the dynamics of polymerized URA7 fibers (Figure 2A). URA7 is a major CTP synthase that catalyzes ATP-dependent conversion of UTP to CTP[37] which, under conditions of starvation, polymerizes out of solution in the cytoplasm into insoluble fibers in a catalytically inactive form. This process is mediated by changes in the surface charge of the protein resulting from changes in cytoplasmic pH, and can be used as an indicator of metabolic state[38, 39]. Under ordinary circumstances, as ATP production increases the vacuolar ATPase pumps more protons out of the cytoplasm which becomes less acidic, leading to URA7 fibers dissociating into soluble active protein (Figure 2B). This is a special case of a more general phenomenon in which cytoplasmic pH and changes in macromolecular crowding affect the physical properties of yeast cytoplasm, with starvation associated with acidic cytosol, increased protein-protein interactions, and a decrease in the function of many metabolic enzymes[40, 41].

**Figure 2.**
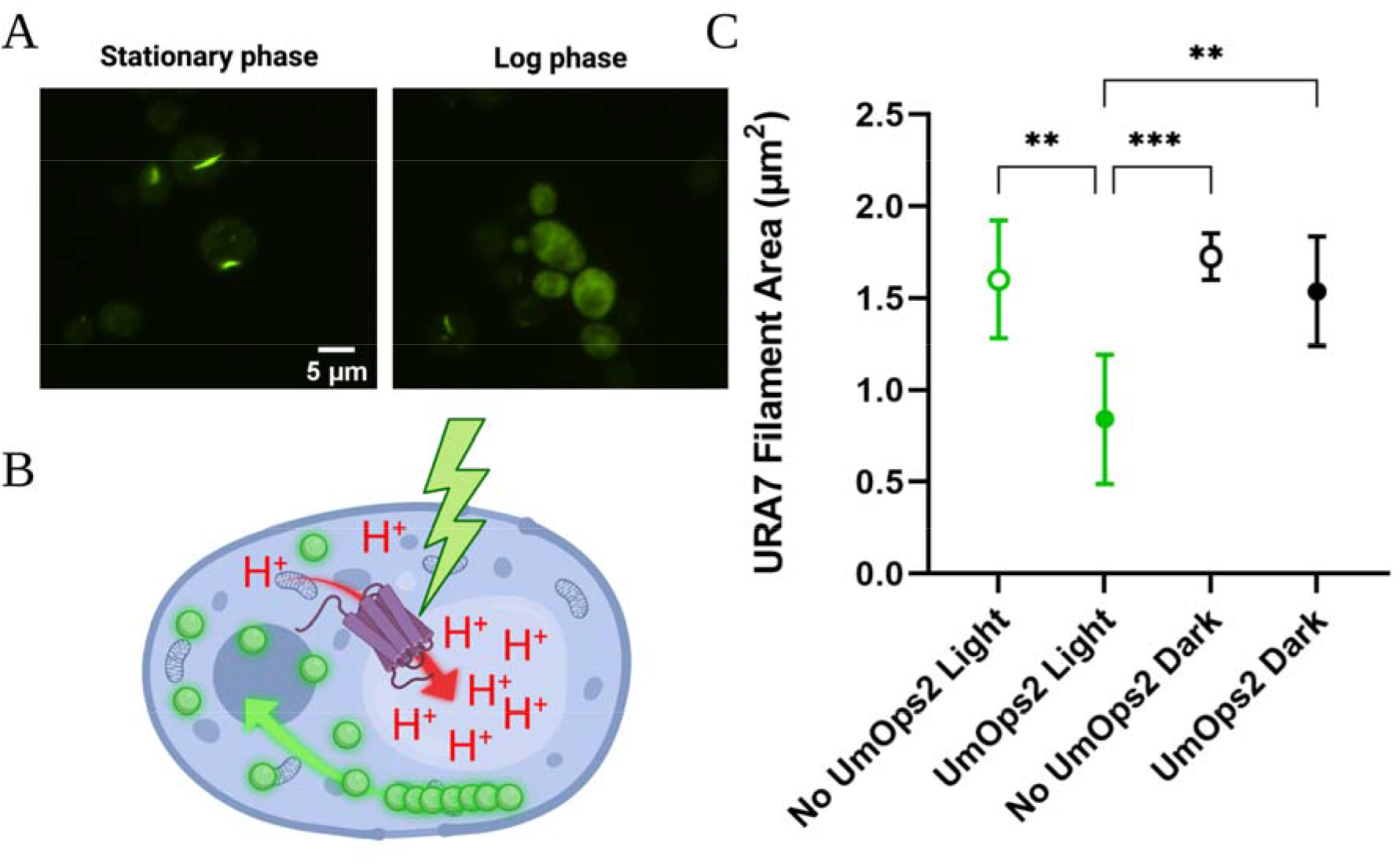
Examining the effect of rhodopsin function on yeast physiology. (A) Yeast bearing URA7-GFP fusion proteins exhibit a single fluorescent fiber of polymerized inactive enzyme in stationary phase, which dissociates into free active enzyme in log phase due to cytoplasmic deacidification. This can be used as a marker of physiological state, with filaments indicating starvation. (B) Model of the effect of vacuolar rhodopsin on yeast physiology. Excitation of rhodopsin with green light pumps protons from the cytoplasm to the vacuole, deacidifying the cytoplasm and dissociating URA7-GFP fibers. (C) Experimental effect of green light on URA7-GFP fiber formation in yeast with and without UmOps2 protein (5 replicates, mean and standard deviation visualized). Cells bearing the vacuolar rhodopsin exhibit significantly smaller URA7 fibers when grown in green light, while those without rhodopsin do not.

We observed this effect of proton pumping on metabolic state by examining GFP-tagged URA7 fibers in light and dark conditions. We grew yeast bearing *URA7-GFP* fusion genes, both with and without UmOps2, for 24 hours in the presence and absence of green LED illumination. Yeast bearing UmOps2 displayed a significant decrease in filament area when cultured under green light, while those without rhodopsins did not (Figure 2C; one-way ANOVA, *F*_3,16_=9.6, *p*=0.0007, pairwise differences assessed with Tukey’s HSD with α=0.05). The reduction in filament area demonstrates a deacidification of the cytoplasm sufficient to alter the function of metabolic enzymes in the presence of green light. While this indicates an effect on cellular physiology and successful pumping of protons across the vacuole membrane, we do not know whether this proton motive force is able to result in ATP synthesis. As a proton pump, rhodopsins are capable of directly deacidifying the cytoplasm even in the absence of ATP production by the vacuolar rotary ATPase. This would still likely result in reduced total ATP consumption as the rotary ATPase becomes inhibited by high proton motive force opposing ATP hydrolysis.

### Fitness consequences of rhodopsin expression

We measured the fitness effect of rhodopsin expression on yeast via a competition assay. Yeast were grown in Yeast Extract Peptone Glycerol media (YEPG) supplemented with all-trans retinal. The purpose of this medium is to force respiration, a state which limits available energy, due to the presence of non-fermentable glycerol as the sole carbon source. When grown in YEPG, yeast cultures typically reach extremely low levels of oxygen and become oxygen-limited, respiring at a fraction of the maximum possible rate their metabolisms could theoretically support[42]. This maximizes the potential for phototrophic energy production to supplement a low rate of respiration in less metabolically active cells. As rhodopsins are known to enhance the survival of bacteria under starvation conditions[17], we allowed cells to experience an extended stationary phase by performing transfers to fresh media only every 48 hours. Yeast cultures were acclimatized to growth in YEPG for 24 hours prior to mixing and competition.

We measured the frequency of rhodopsin-bearing cells versus control cells after mixing them at a 1:1 ratio, and calculated their apparent selective advantage in dark and light conditions (see Methods) after 72 hours of culture (Figure 3). Our control strain was isogenic, but expressing GFP under the same promoter as the rhodopsin. Rhodopsin-bearing yeast experienced increased fitness by a factor of 2% in light relative to dark, as calculated using the ratio of Malthusian growth parameters[43–45] (two-tailed *t*-test, *t*=3.35, *n*=10, *p*=0.017).

**Figure 3.**
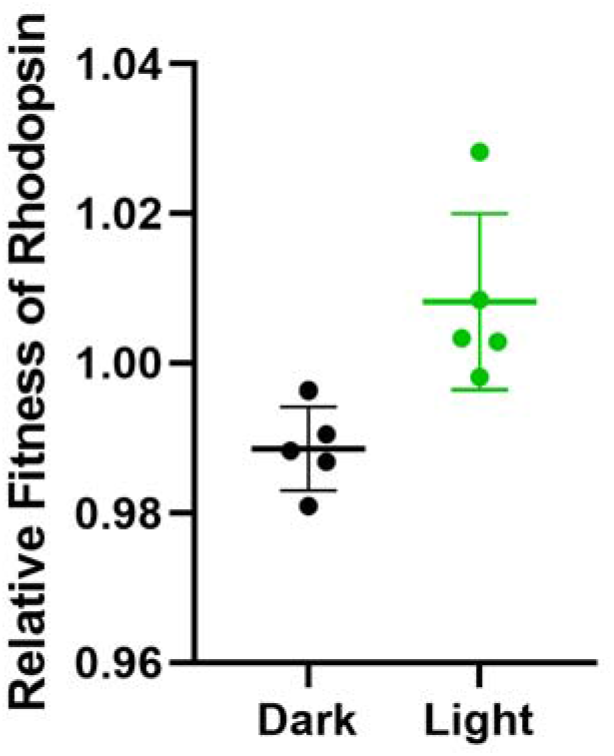
Selective advantage of rhodopsin-bearing yeast. Control yeast (GFP, under the same promoter as the rhodopsin strain) and yeast bearing UmOps2 were grown together in dark and light conditions for 2 passages. Rhodopsin-bearing yeast were on average 2.2% more fit, measured as the ratio of Malthusian growth parameters, when grown in green light compared to when they were grown in the dark.

### Cell viability

Fundamentally, there are only two ways that a replicator can increase in frequency in a population: it can reproduce more, or die less[46]. While rhodopsin-bearing yeast grown in light have an advantage during competition (Figure 3), it was not clear whether this benefit stems from greater reproduction or reduced mortality, which we expect would be most impactful during the extended starvation experienced in stationary phase. To disentangle these mechanisms, we measured the viability of cells from each strain during extended stationary phase culture. We inoculated GFP (control) and UmOps2-bearing yeast into YEPG supplemented with all-trans-retinal, with or without green light. We allowed them to grow for 72 hours without changing the growth medium. Every 24 hours, we extracted yeast, measuring viability with the live/dead cell stain propidium iodide (Figure 4). Overall cell death across all populations was relatively low, ≤3%, and yeast incubated at stationary phase accumulated dead cells at or slower than ~0.3% per 24 h. Yeast bearing rhodopsins unexpectedly experienced 78% higher mortality than GFP controls rather than decreased mortality across growth conditions, while culture under continuous green light increased mortality by 282% (ANOVA summary in Table S1, all parameter estimates significant at p<0.001). This is consistent with observations that yeast bearing rhodopsins experienced a 1.1% fitness deficit in the dark compared to control yeast (two-tailed *t*-test, *t*=-4.5, *p*=0.0105) (Figure 3). Taken together, this indicates that the selective advantage experienced by rhodopsin-bearing yeast under illumination is due to increased growth, which is sufficient to overcome their measured increased mortality.

**Figure 4.**
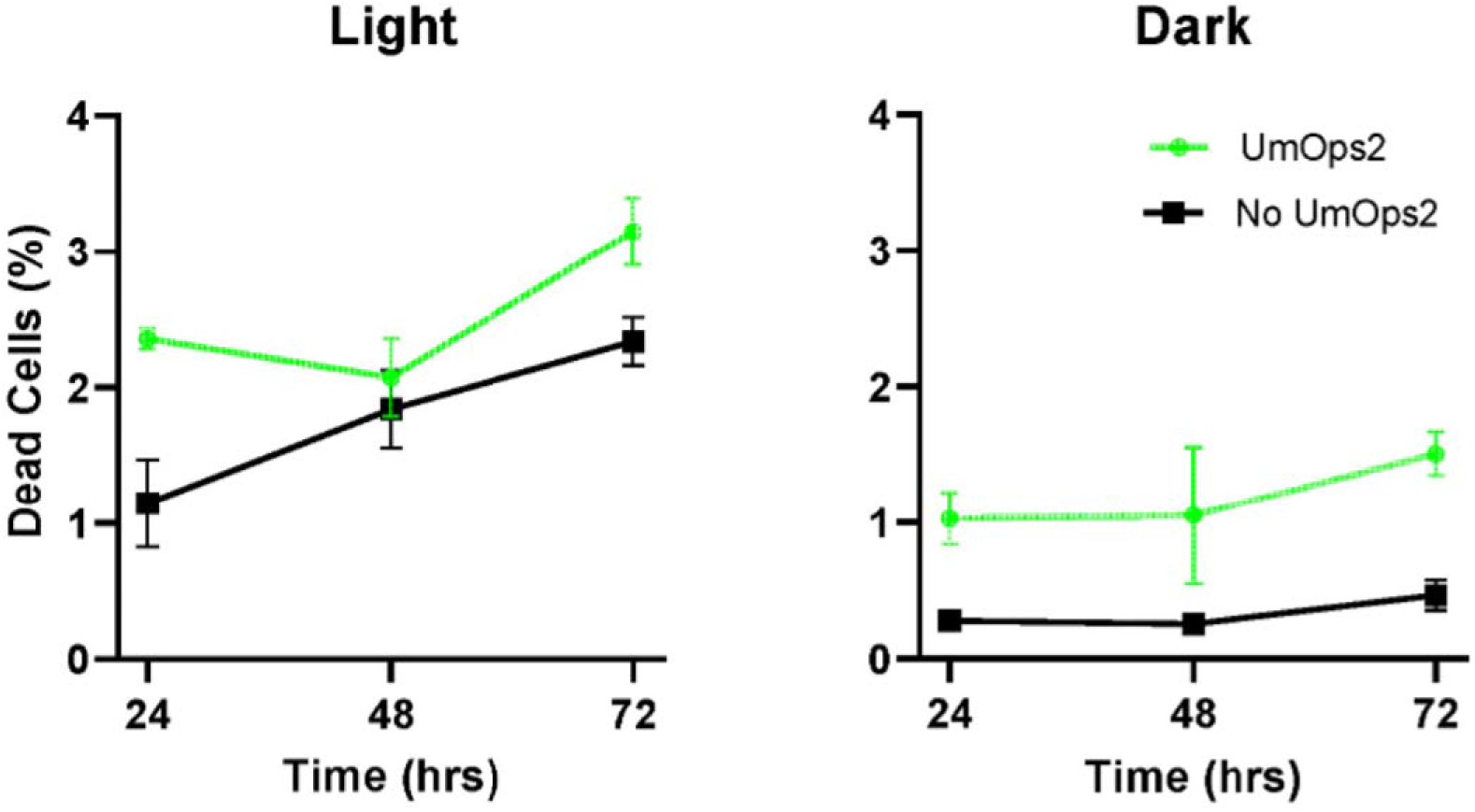
Effect of rhodopsin on cell viability in stationary phase. Control and UmOps2-bearing yeast were grown in YEPG for 3 days in dark and light conditions, stained daily with propidium iodide to measure cell death in each population. Each datapoint in represents the mean of 5 replicate UmOps2+ and UmOps2-populations ± one standard deviation Rhodopsin increased cellular mortality in both light and dark, suggesting that the higher fitness of rhodopsin-bearing yeast during competition (Figure 3) is due to increased growth, not reduced death.

## Discussion

In this paper, we explore the capacity of a heterotrophic eukaryote with no known history of phototrophy, *S. cerevisiae*, to become a facultative phototroph. We transformed *S. cerevisiae* with a fungal rhodopsin, *UmOps2*, from the corn pathogen *Ustilago maydis*. UmOps2 protein localized to the yeast vacuole and proved physiologically functional, reducing indications of starvation and quiescence under light, with no impact on physiology in the dark. Under light, UmOps2 increased fitness by 2%, which was due entirely to increased growth rate rather than reduced cellular death in stationary phase.

The success of this artificial horizontal transfer of rhodopsin from one fungal species to another underscores what has been learned from phylogenetic reconstructions: rhodopsins, as physiologically flexible genes not dependent upon a specific organismal context to provide a selective advantage, are easily capable of functional horizontal transfer, even in eukaryotic cells with complex architectures. Previous attempts to transfer proton-pumping rhodopsins to eukaryotic species in a way that contributes to cellular bioenergetics have generally targeted the mitochondrion[31–34], the eukaryotic membrane most classically associated with ATP production, even though eukaryotic proton-pumping rhodopsins typically localize to the cell membrane, vacuole, or other endomembrane system components[28, 47, 48]. These mitochondrial localizations have always involved fusion of the rhodopsin sequence to a mitochondrial inner membrane protein[31–34]. This suggests that natural eukaryotic proton-pumping rhodopsins do not localize to the mitochondrion because multiple evolutionary steps are needed in tandem for this to occur from the starting point of a horizontally transferred rhodopsin. Without specific mitochondrial localization signals, membrane proteins are typically integrated into the endomembrane system, particularly the endoplasmic reticulum.

It has recently been shown that eukaryotes can obtain useful work from rhodopsins localized to the endoplasmic reticulum – diatoms bear a proton-pumping rhodopsin specifically localized to the Chloroplast Endoplasmic Reticulum Membrane (CERM) that encloses their secondary plastid, which is believed to enhance the efficiency of their carbon-concentration mechanism[48] in order to reduce necessary RuBisCo protein for carbon fixation. A functional role in the energetics of the endomembrane system of a eukaryote could represent an evolutionary ‘foot in the door’, allowing maintenance of a gene without more specialized localization signals. Vacuolar localization for more specialized phototrophic energy production, requiring point mutations targeting trafficking via the ALP pathway[49], would then be free to evolve gradually rather than via dramatic fortuitous gene fusions. Thus, targeting to the vacuole may simply be more evolutionarily accessible than targeting to the mitochondrial inner membrane, and these vacuolar rhodopsins would then be free to spread further through the eukaryotes via additional horizontal gene transfer.

While the function of UmOps2 was dependent upon supplementation with all-trans retinal in our experiment, this may not represent a barrier to horizontal transfer of a complete rhodopsin system. Rhodopsins are known to occasionally transfer in multigene cassettes bearing a retinal synthesis pathway[12]. Organisms with proton-pumping rhodopsins that lack retinal-production pathways are also known; they typically acquire the pigment by consuming rhodopsin-bearing bacteria[50].

Rhodopsin expression carries a cost in our model system. While exhibiting a selective advantage in the presence of green light compared to dark, rhodopsin-bearing yeast face a selective disadvantage in the dark, and consistently exhibit a higher rate of cell death in both light and dark conditions. The reasons for this cost are unclear. It is unlikely to be a result of sheer protein biosynthesis costs, as this rhodopsin and the GFP expressed in control cells are of similar mass (27 kilodaltons for GFP versus 32 kilodaltons for UmOps2), and are driven by the same promoter. However, whatever the source of this fitness cost, it is apparently less than the growth advantage provided by the ability to acquire energy as a photoheterotroph in the light.

UmOps2 represents one of three described classes of fungal rhodopsins in ascomycetes.

It is an LR-like rhodopsin, which function as rapid proton-pumps, as compared to CarO-like and NR-like rhodopsins, which function as sensory proteins[51]. Notably, both the prototypical rapid proton pumping rhodopsin *LR* from the fungus *Leptosphaeria maculans* and the LR-type rhodopsin UmOps2 from *Ustilago maydis[28]* localize primarily to the vacuole membrane when expressed in *S. cerevisiae* (data not shown). Our results demonstrating that UmOps2 is capable of pumping physiologically relevant quantities of protons into the vacuole suggests that energy production via eukaryotic vacuolar light-driven proton pumps could be a general pattern, with the vacuole being a eukaryotic membrane where proton pumping is particularly useful. Here, this activity can alleviate the ATP cost of acidification of the vacuole/endosome compartment or raise the membrane potential high enough to run the rotary ATPase in reverse, either way transducing light into biologically-available energy. As a bioenergetically-relevant membrane, this compartment could be a common location of rhodopsins in photoheterotrophic eukaryotes.

Artificial photoheterotrophic metabolism may have practical applications. *S. cerevisiae* is frequently used as an industrial organism for biofuel and bioproduct production applications[52]. Photoheterotrophic metabolism stands to potentially increase their yield, as more substrate might be able to be put towards production of the desired product instead of towards catabolism and energy metabolism. The success of this rerouting of substrate towards product production would depend on the relative energy flux per unit protein of rhodopsins versus glycolytic enzymes and respiratory electron transport chains, as has been previously examined in studies of ‘overflow metabolism’ in bacteria[53]. Exploration of this could be of interest for biofuel or bioproduct production in environments with limited material availability but high potential light availability, such as the industrial production of secondary modified metabolites or perhaps even aerospace applications.

The successful transfer of photoheterotrophic metabolism to a well-understood model organism via simple synthetic biology techniques may also open novel research avenues. The origin of phototrophy is one of the most impactful innovations in the history of life on Earth, and it is likely that rhodopsin-based retinalophototrophy predates chlorophototrophy as its earliest form[54]. Recent work has reconstructed ancient ancestors of modern rhodopsins, and probed their likely function[55, 56]. While rhodopsins have been successfully expressed in bacteria before[26, 57], a cellular environment likely more akin to the prokaryotic cells of the early Earth, many eukaryotes bear these proteins and may have done so from the origins of eukaryotes themselves[16, 58, 59]. The ability to easily synthesize, express, and characterize the function of ancient rhodopsins in S*. cerevisiae*, a workhorse of modern cell biology, opens a range of new opportunities for understanding the origins and subsequent evolution of retinalophototrophy.

In summary, we transformed the yeast *Saccharomyces cerevisiae* into a facultative photoheterotroph by adding a single vacuole-localized rhodopsin. When exposed to green light, rhodopsin-bearing yeast exhibited a modest but significant fitness advantage. This required no evolutionary optimization, illustrating the ease of horizontal transfer of this phototrophic system and potentially explaining the repeated gain of rhodopsin-mediated phototrophy in eukaryotes. Using synthetic biology to construct photoheterotrophic organisms could have benefits for industrial purposes, as it is another method to provide energy for cellular metabolism over and above chemical feedstocks. Synthetic construction of photoheterotrophic yeast provides novel insights into the evolution and spread of retinalophototrophy, its optimization and integration into previously heterotrophic metabolisms, and stands to open new avenues of research and experimentation.

## Methods

### Engineering & Strains

The *U. maydis* rhodopsin *UmOps2* reading frame was codon optimized and synthesized via ThermoFisher, as was *GFP*, flanked by restriction sites SpeI and AflII. Both *GFP* and *U. maydis* rhodopsin UmOps2 were ligated into a custom expression vector driven by a *TEF1* promoter and *CYC1* terminator, associated with a *NAT-NT2* resistance cassette and a fragment of *HO* gene homology bearing the loss-of-function mutations present in S288C. By cutting within this *HO* homology using AfeI or BsaAI, *UmOps2* or *GFP* can be introduced as a high-copy repeat at the *HO* locus.

*UmOps2* and *GFP* were introduced at high copy to the *HO* locus of homozygous diploid Y55 derivative Y55HD previously used by the Ratcliff laboratory[60]. These strains were sporulated, and the resultant haploid heterothallic spores mated with each other to produce homozygous diploid control yeast homozygous for *GFP* at the *HO* locus and experimental yeast homozygous for *UmOps2 at* the *HO* locus.

*GFP* was fused to *URA7* at the C-terminus via PCR transformation from plasmid pYM25[61] bearing *GFP* and *hphNT2* in the Y55HD genetic background[60]. Heterozygous yeast were used as a control for metabolic analysis, and after mating with yeast bearing *UmOps2*, double heterozygotes were used as experimental strains for metabolic analysis of the effect of rhodopsin on URA7 fiber morphology.

### Cell Culture and Competition

All yeast were grown in liquid Yeast Extract Peptone Glycerol (YEPG) media supplemented with 10 microliters of 10 mM all-trans retinal in 100% ethanol, for a final concentration of 10 μM of all-trans retinal. This supplemented medium was used for all steps.

For competition experiments, control and rhodopsin bearing yeast were grown separately to saturation shaking at 250 RPM at 30 °C for 24 hours. On Day 0, 500 μL of both GFP-bearing controls and rhodopsin-bearing experimental cells were mixed. This mixture was measured via flow cytometry to determine the baseline ratio of control and rhodopsin-bearing yeast. A starting culture of 50 μL of mixed stationary-phase culture was transferred two new tubes of retinal-supplemented YPG. One was placed in an ordinary shaking incubator at 250 RPM and 30 °C, while another was placed in another shaking incubator with the same settings supplemented with green LEDs and a reflective foil wrapping to provide bright green illumination, measured at an average intensity of 1996 lux. These cells were allowed to grow for 48 hours, upon which 50 μL of stationary phase culture was transferred to new tubes and placed into the same incubator. At 24 and 72 hours, samples were taken for flow cytometry to count GFP and rhodopsin bearing yeast.

### Fitness quantification

Frequency of different cell types were determined either via flow cytometry or microscopy, as described above. Relative fitness can be determined from the change in the frequency genotypes over time. Relative fitness was determined as described in the Lenski long-term evolution experiment[44, 45]. The relative fitness *W* of a rhodopsin-bearing yeast can be defined as the ratio of the number of generations of rhodopsin-bearing yeast (*G_R_*) divided by the number of generations of a control yeast (*G_C_*) over the same time frame using Equation 1.

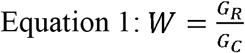

The number of generations experienced by a genotype in a population (*G*) after a number of transfers *n* during which the population reaches saturation can be calculated from the fraction of the population at passage (*F_n_*), the fraction of the population at passage 0 (*F*_0_), the dilution factor (*d*), and the number of transfers (*n*) using Equation 2:

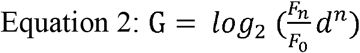

By combining Equation 1 and Equation 2, we determine the equation for relative fitness *W* in terms of number of passages (*n*), fraction of the population bearing rhodopsin after passage *n* (*Fr_n_*) and at passage 0 (*Fr*_0_), fraction of the population that are controls after passage *n* (*F_C_n__*) and at passage 0 (*F*_*C*_0__), the dilution factor (*d*), in the form of Equation 3:

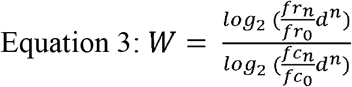

Equation 3 was used to calculate the relative fitness of rhodopsin-bearing yeast in dark and light incubation for all 5 replicate populations each at day 3 of cultivation, such that dilution factor *d* equals 200 and number of passages *n* equals 2.

### Cell Viability

In order to quantify the viability of cells, a propidium iodide stain was used to measure the proportion of dead cells as a rhodopsin-bearing population and control population progressed through stationary phase. Five replicate populations were inoculated and grown in YP Glycerol supplemented with all-trans retinal for 72 hours. After 24, 48, and 72 hours, 500 μL of each population was spun down and double washed, resuspending in 1 mL of water for imaging. One μL of propidium iodide stock solution (1 mg/mL) was added to the pellet and was imaged on a TI Nikon Eclipse Ti-Motorized Inverted Microscope. Four fields of view were imaged at 20x objective magnification, capturing a bright-field image, and an image with TRITC filter and 1.3 ms exposure to capture propidium iodide stained dead cells. Images were thresholded using MaxEntropy auto thresholding in Fiji Image Analysis to identify and count dead cells, and the bright-field channel segmented via Cellpose[62] to count total cells.

### URA7 Filament Metabolic Analysis

In order to determine whether rhodopsin provides an energy benefit, metabolic state was assessed via polymerization of GFP-tagged URA7p. A starting culture of 50 μL of control yeast bearing GFP-tagged URA7p, and experimental yeast bearing this fusion and the UmOps2 rhodopsin, were inoculated and grown for 24 hours in 10 mL Synthetic Complete media supplemented with all-trans retinal. Five replicates each were grown in a dark incubator at 250 RPM and 30 °C, and a green illuminated incubator at 30 °C with light approximately 1996 lux. At 24 hours growth, stationary phase cells were removed and quickly imaged on a TI Nikon Eclipse Ti-Motorized Inverted Microscope. Fibers were captured in the FITC channel with a constant exposure of 1s and analog gain of 64.0x. The average filament area was quantified using Fiji Image Analysis via automatic image segmentation using MaxEntropy auto thresholding of particles ≥ 10 pixels in size.

#### Strains used

yAB1: Y55HD background, *URA7/URA7-GFP:HphNT2*
yAB406: Y55HD background, *ho:UmOps2:NatMX6/ho:UmOps2:NatMX6*
yAB414: Y55HD background, *ho:UmOps2-GFP:NatMX6/ho:UmOps2-GFP:NatMX6*
yAB452: Y55HD background, *URA7/URA7-GFP:HphNT2 HO/ho:UmOps2:NatMX6*
yAB703: Y55HD background, *ho:GFP:NatMX6/ho:GFP:NatMX6*

#### Plasmids used

pYM25: *GFP* for protein fusion and a *hphNT2* resistance cassette[61]
pWR112: *UmOps2* under a *TEF1* promoter, *NATMX6* resistance, *HO* homology for chromosomal insertion
pWR113: Expression of *UmOps2-GFP* under a *TEF1* promoter, *NATMX6* resistance, *HO* homology for chromosomal insertion
pWR162: Expression of *GFP* under a *TEF1* promoter, *NATMX6* resistance, *HO* homology for chromosomal insertion

#### Oligonucleotides used for URA7-GFP fusion

Ura7_S2: TAGAAATTTATCTTTGTTAATGCAGTAGACTTTTAATTCTAAAATTTTGATTTAATCG ATGAATTCGAGCTCG
Ura7_S3: GTCATTGAAGGTAAGTACGATCTTGAGGCCGGCGAAAACAAATTCAACTTTCGTACG CTGCAGGTCGAC

## Acknowledgements

Figures 1C and 2B created using BioRender.com.

## Author Contributions

AB conceived of the research. AP, AB and WCR planned the experiments. AP and AB performed the experiments and analyzed the data. All authors contributed to writing.

## Research support

This work was supported by a NSF Graduate Research Fellowship to AP, and NSF grant DEB-1845363, Human Frontiers Grant RGY0080/2020 and a Packard Foundation Fellowship for Science and Engineering to WCR.

## Supplementary Tables

**Table S1.** Impact of rhodopsins, culture in light vs. dark, and time in stationary phase via ANOVA. Dependent variable: cell viability. Independent variables: genotype (rhodopsin vs. GFP control), culture conditions (light vs. dark), and time (24, 48, 72h). As we did not see any clear sign of GxE interactions in our experiment, we did not examine these interactions statistically. For this ANOVA, F_3,59_ = 48.9, p<0.0001, and the adjusted r^2^ =.71.

| Analysis of Variance |  |  |  |  |
| --- | --- | --- | --- | --- |
| Source | DF | Sum of Squares | Mean Square | F Ratio |
| Model | 3 | 0.0045 | 0.0015 | 48.87 |
| Error | 56 | 0.0017 | 0.000031 | **Prob > F** |
| C. Total | 59 | 0.0062 |  | <.0001* |

Table S1. **Impact of rhodopsins, culture in light vs.**
| Parameter Estimates |  |  |  |  |
| --- | --- | --- | --- | --- |
| Term | Estimate | Std Error | t Ratio | Prob>\|t\| |
| Intercept | 0.0088 | 0.0019 | 4.67 | <.0001* |
| Rhodopsin vs. Control | -0.0042 | 0.00071 | -5.86 | <.0001* |
| Light vs. Dark | -0.0071 | 0.00071 | -10.00 | <.0001* |
| Time | 0.00013 | 3.6e-5 | 3.51 | 0.0009* |

